# Gene expression QTL mapping in stimulated iPSC-derived macrophages provides insights into common complex diseases

**DOI:** 10.1101/2023.05.29.542425

**Authors:** Nikolaos I Panousis, Omar El Garwany, Andrew Knights, Jesse Cheruiyot Rop, Natsuhiko Kumasaka, Maria Imaz, Lorena Boquete Vilarino, Anthi Tsingene, Alice Barnett, Celine Gomez, Carl A. Anderson, Daniel J. Gaffney

**Author notes:** These authors jointly supervised this work.

## Abstract

Many disease-associated variants are thought to be regulatory but are not present in existing catalogues of expression quantitative trait loci (eQTL). We hypothesise that these variants may regulate expression in specific biological contexts, such as stimulated immune cells. Here, we used human iPSC-derived macrophages to map eQTLs across 24 cellular conditions. We found that 76% of eQTLs detected in at least one stimulated condition were also found in naive cells. The percentage of response eQTLs (reQTLs) varied widely across conditions (3.7% - 28.4%), with reQTLs specific to a single condition being rare (1.11%). Despite their relative rarity, reQTLs were overrepresented among disease-colocalizing eQTLs. We nominated an additional 21.7% of disease effector genes at GWAS loci via colocalization of reQTLs, with 38.6% of these not found in the Genotype–Tissue Expression (GTEx) catalogue. Our study highlights the diversity of genetic effects on expression and demonstrates how condition-specific regulatory variation can enhance our understanding of common disease risk alleles.

## Introduction

Disease-associated variants are often located in noncoding regions of the genome ^1, 2^, making biological interpretation of their function challenging. If these variants alter gene expression this should, in principle, be detectable by mapping expression quantitative trait loci (eQTL). eQTLs have now been discovered across a broad range of human tissues and cell types ^3, 4^, with at least one eQTL known for every human gene. Despite this, a surprisingly large fraction of disease associations do not share a causal variant with a known eQTL ^5–8^. For example, the most recent publication by the Genotype-Tissue Expression (GTEx) consortium, which mapped eQTLs across 49 tissues ascertained from 838 individuals, reported that approximately 43% of disease associations colocalize with a detectable eQTL ^3^.

One potential explanation for missing disease-associated eQTLs is that many regulatory variants may function in highly specific cellular contexts, such as activated immune cells. To address this, multiple studies have now been performed using bulk RNA-sequencing of activated cell conditions ^9–17^ and more recently using single cell approaches ^18, 19^. However, because primary cell material is frequently limited, eQTL mapping has typically been restricted to a relatively small set of environmental contexts following perturbation with a limited set of classical stimuli.

Here, we differentiated induced pluripotent stem cells (iPSCs) from 209 individuals to macrophages and mapped eQTLs across twelve different cellular conditions at two timepoints post stimulation. We used this dataset (MacroMap) to explore the properties of response eQTLs (reQTLs) and their relevance for disease.

## Results

### Expression profiling of macrophage innate immune responses

We selected 217 iPSC lines derived from unrelated healthy donors by the HipSci consortium ^20^, and differentiated them to macrophages using a previously described protocol ^15^ with minor modifications (see Methods and Supplementary note). During the differentiation process we collected macrophage precursor cells at day 0 (labelled “Prec_D0”) and day 2 (“Prec_D2”) and profiled their transcriptome via a low-input RNA-seq protocol. We also profiled the transcriptome of unstimulated iPSC-derived macrophages after six and 24 hours. We next perturbed the cells with a panel of ten different stimuli and profiled gene expression six and 24 hours after stimulation (Figure 1a). Our stimulation panel (Supplementary table 1) included stimuli that trigger pro- and anti-inflammatory pathways in macrophages (IFNβ, IFNγ, interleukin-4/IL4) and those that induce response to viral infection (Resiquimod/R848, Poly I:C/PIC). We also included combinations of stimuli to mimic bacterial response to infection (smooth lipopolysaccharide/sLPS, Pam3CSK4/P3C, CD40 ligand + IFNγ + sLPS/CIL, interleukin-10 + sLPS/LIL10). Furthermore, we stimulated macrophages with myelin basic protein (MBP) to mimic the innate immune response of microglia (brain-resident macrophages) to stimuli. For simplicity throughout, we include day 0 and day 2 precursor cells when we refer to “stimulated conditions’’, except where otherwise stated. We profiled the transcriptome of 5,208 samples and after quality control (Methods) retained 4,698 unique RNA-seq libraries from 209 unique iPSC lines (Supplementary figure 1).

**Figure 1:**
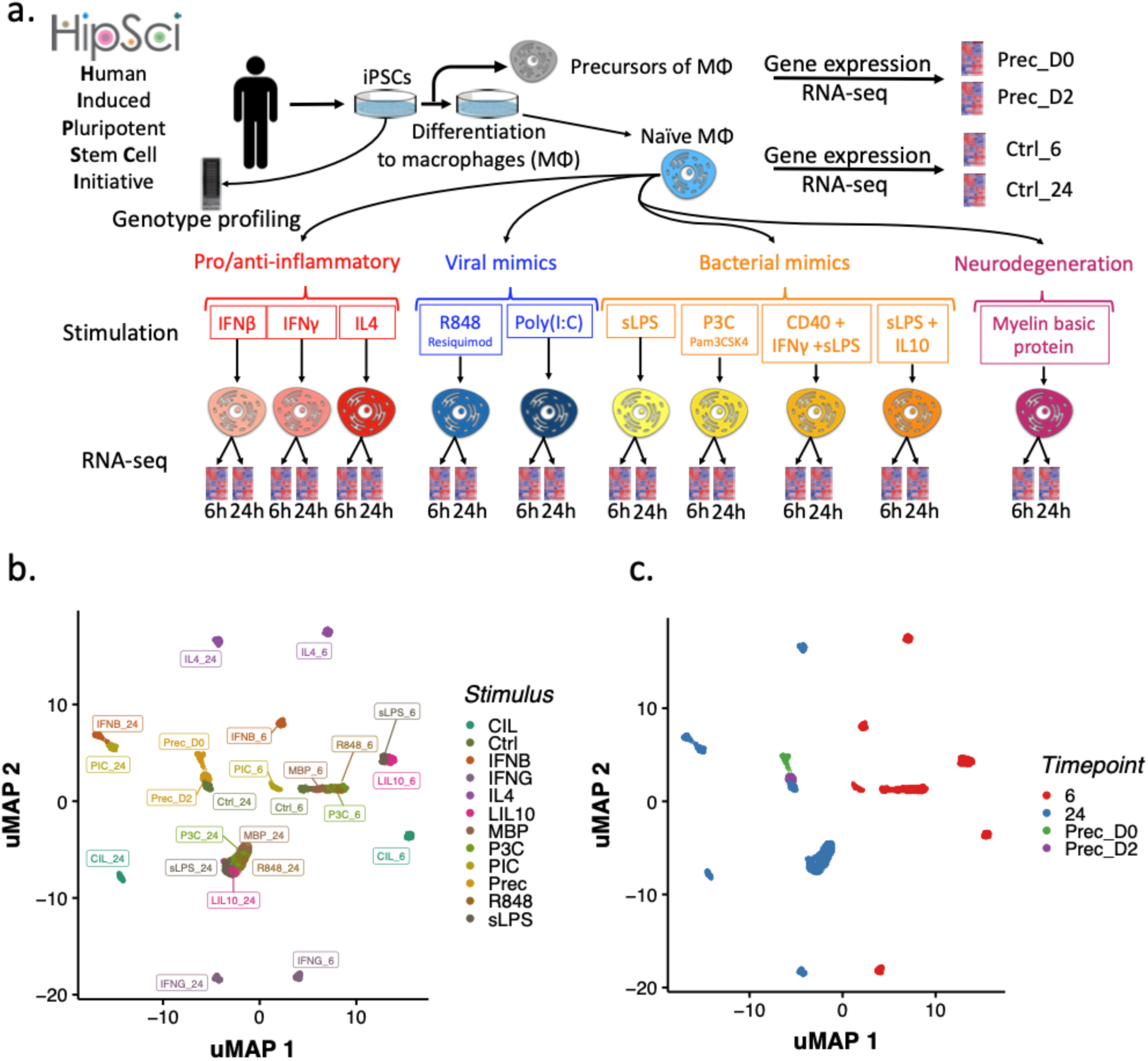
Experimental Workflow and Transcriptomic Analysis. **a.** Overview of the experimental workflow. Isolation and stimulation of naive macrophages and measurement of gene expression via RNA-seq at two post-stimulation time points (6h and 24h) for multiple stimuli. **b/c** UMAP representation of 4698 RNA-seq libraries after quality control (Methods) coloured by different stimulation conditions (b) and time point (c).

We first explored which technical and biological confounders affected gene expression using variance component analysis (Supplementary figure 1d) and identified stimulation as the second most important driver of the total expression variation (29.3%) after the library preparation method (39.9%) (Supplementary figure 1e).

A UMAP projection of the gene expression data showed that samples clustered by stimulation and time point (Figure 1b, c). To better quantify the structure present in the gene expression data we estimated the pairwise Pearson’s correlation (r) between conditions (mean TPM values per gene across all individuals for a given condition) (Supplementary Figure 2a). For example, the gene expression profile of precursor cells at day 2 clustered closely with samples from naive cells at the 24 hour time point (Pearson r=0.98). Likewise, gene expression in macrophages stimulated with PIC was highly correlated with gene expression in IFNβ at both time points, highlighting an expected overlap in signalling because PIC increases IFNβ expression ^21^.

We next identified differentially expressed genes (DEGs) between naive and stimulated conditions using DESeq2 ^22^ (Methods). On average, we found a median of 2,306 DEGs (false discovery rate (FDR) = 5%, fold change ≥2) between naive and stimulated cells across all conditions (Supplementary Tables 2,3,4), although the number varied widely between conditions (488-5,427 DEGs) (Supplementary Figure 2b). These DEGs were enriched in gene ontology (GO) terms (Supplementary Table 5-6) and Reactome pathways ^23^ (Supplementary Tables 7-8) corresponding to relevant immunological pathways. For example, “response to bacterium” (GO:0009617) was the most enriched GO term in DEGs six hours after sLPS stimulation, with *TNF* upregulated 13.75-fold ^24^. Likewise, “Response to virus” (GO:0009615) was the GO term most enriched with genes differentially expressed in cells six hours after stimulation with IFNβ, with *GBP1* (Guanylate Binding Protein 1) upregulated 256-fold ^25–27^. This demonstrates that iPSC derived macrophages can faithfully recapitulate the known biological pathways activated by different stimuli.

### Genetic regulatory effects across different stimulation conditions

To investigate genetic regulation of gene expression we mapped expression quantitative loci (eQTL) across all twenty four conditions (Methods). We mapped cis-eQTLs within ±1Mbp of the transcription start site (TSS) of each gene at a false discovery rate (FDR) of 5%. We found between 1,781-3,735 eGenes (genes with at least one cis-SNP significantly associated with expression) per condition (Supplementary Figure 3a), with an enrichment of eQTL lead SNPs (eSNPs) in close proximity to promoters (P=1.2×10^-124^, mean log odds ratio= 2.29) (Supplementary Figure 3b). We performed conditional eQTL mapping to identify additional cis-eQTLs with independent effects on expression (Methods) and found that 3.3% - 8% of eGenes in each of our 24 cell conditions, and 18% of all eGenes, had more than one eSNP (1817 eGenes) (Supplementary Figure 4a). Secondary and tertiary eSNPs (conditionally independent eQTLs) were further away from the TSS (mean distance 206 and 233 kb respectively) compared to the primary signals (mean distance 94 kb Wilcoxon p=1.1×10^-215^, p=5.7×10^-16^ respectively) (Supplementary Figure 4b).

Per condition, our eQTL detection rate (22.4% of tested genes, mean over all conditions) was lower compared to GTEx tissues of similar sample size (35% in GTEx tissues of between 180-210 individuals). One likely explanation for our lower detection rate is that we studied a single cell type, while most GTEx eQTL studies are derived from complex tissues. Consistent with this, GTEx eQTLs in lymphoblastoid cell lines (LCLs), had an equivalent detection rate to our study (∼22%). Summing over all cell conditions we identified 10,170 unique eGenes (72.4% of expressed genes) (Figure 2a). This is higher than expected given our study sample size (mean across all conditions n=202), even compared to studies of complex tissues. For example, the number of eGenes discovered in GTEx liver (n=208) and brain cortex (n=205) was 4415 and 7108, equivalent to 25.6% and 37.5% of expressed genes, respectively. Our overall eQTL detection rate also exceeds that observed for individual cell types studied in GTEx. For example, we detected a similar proportion of eQTLs to that found in GTEx cultured fibroblasts (12,280 eGenes, 73.2% of expressed genes), a study with over double our sample size (n=483). Although studying many conditions likely revealed additional eQTLs, the large number of eQTLs we detected is likely also driven by the very high degree of biological replication in our study. We note that even when we compared our control conditions at six and 24 hour timepoints, we detected an additional 774 new eGenes (Supplementary Figure 4c). Thus, generating additional gene expression data from the same individuals increased our power to detect eQTLs relative to other studies of similar sample size.

**Figure 2.**
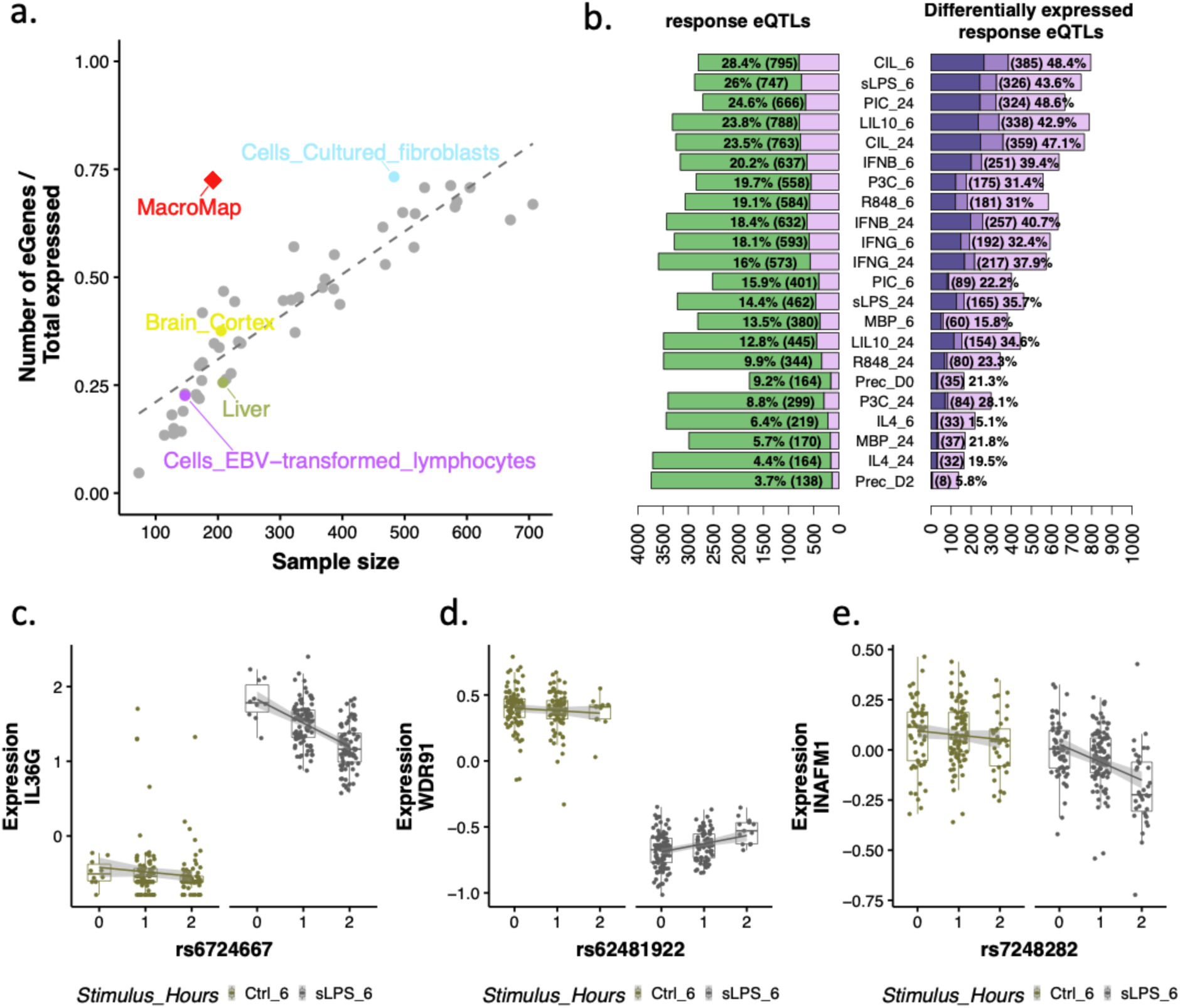
Stimulation-induced changes in gene expression: identification and characterization of reGenes in multiple conditions. **a.** Fraction of expressed genes (protein coding and lincRNAs) that are eGenes (genes with at least one eQTL) in our study (red diamond) and in different GTEx tissues (circles) as a function of the sample size (mean sample size across all conditions in our study). GTEx studies of single cell types (cultured fibroblast and lymphocytes) and tissues with similar sample size to our study (Brain-cortex and Liver) are highlighted. **b.** Number and fraction of reGenes (pink) and the total number of eGenes (green) per stimulation condition (left hand panel) and number and fraction of reGenes that were differentially expressed (dark purple, up-regulated, dark pink, down-regulated) and not differentially expressed (pink) (right hand panel). Examples of genes where reQTL was detected following stimulation and genes were significantly up- (**c),** downregulated (**d**) or where there was no significant change in expression (**e**) in the stimulated condition.

### Stimulation specific genetic regulation of gene expression in immune response

To better establish which cis-eQTLs were truly restricted to stimulated cells, we used mashr ^28^, which compares eQTL effect size estimates between conditions. Using the “common baseline” mode (Methods), mashr estimates the extent to which the eQTL effect size in each stimulated condition deviates from a baseline condition, here defined as the Ctrl_24 condition. Mashr produces a local false sign rate (lfsr) that measures the confidence in the direction of each genetic effect compared to the baseline effect ^29^.

We defined a response eQTL (reQTL) as a significant difference (lfsr<0.05) in genetic effect between the baseline condition and at least one stimulated condition. The number of reQTLs we detected varied widely between conditions, from 3.7% of all eQTLs in precursor cells at day 2 (Prec_D2) to 28.4% of all eQTLs in cells stimulated with CIL at the 6h time point (Figure 2b). Across all conditions, 23.4% (2378) of eGenes had a reQTL in at least one condition with the majority of these (21.9%, 2228) having a larger effect following stimulation than in the naive condition. Approximately 9% (159) of conditionally independent eQTLs were classified as reQTLs. 89% (142) of these genes with a conditionally independent reQTL also had a primary reQTL, an almost 4-fold enrichment (Fisher’s exact test p-value=3^-36^, OR=12.93, 95% CI 7.7 - 23.05) (Supplementary Figure 5b).

Our ability to detect eQTLs for lowly expressed genes is limited. Many of our reQTLs may therefore have similar effect sizes in both the naive and stimulated conditions, but these effects can only be detected as different once gene expression increases in response to stimulation. We found that between 5.8%-48.4% of reQTL genes were differentially expressed (example signals Figure 2c,d) (FDR 5%, fold changeB*) (31.2% on average across conditions), the majority of which (mean across conditions 75.8%) were up-regulated (Figure 2c) compared to the naive condition. Nonetheless, the majority of reQTLs we identified (51.6%-94.2%) were in genes where we were unable to detect a substantial change in expression between the naive and stimulated conditions based on our chosen criteria (Figure 2e).

Next, we asked how widely shared reQTLs were between stimulated conditions, using mashr. Very few (1.1%) reQTLs were specific to a single condition, 90.5% of reQTLs were shared in five or more conditions, with 49% detected in at least half of the conditions (Figure 3a, b). This suggests that the majority of the reQTLs we identified could have been detected with a smaller set of stimulated states.

**Figure 3:**
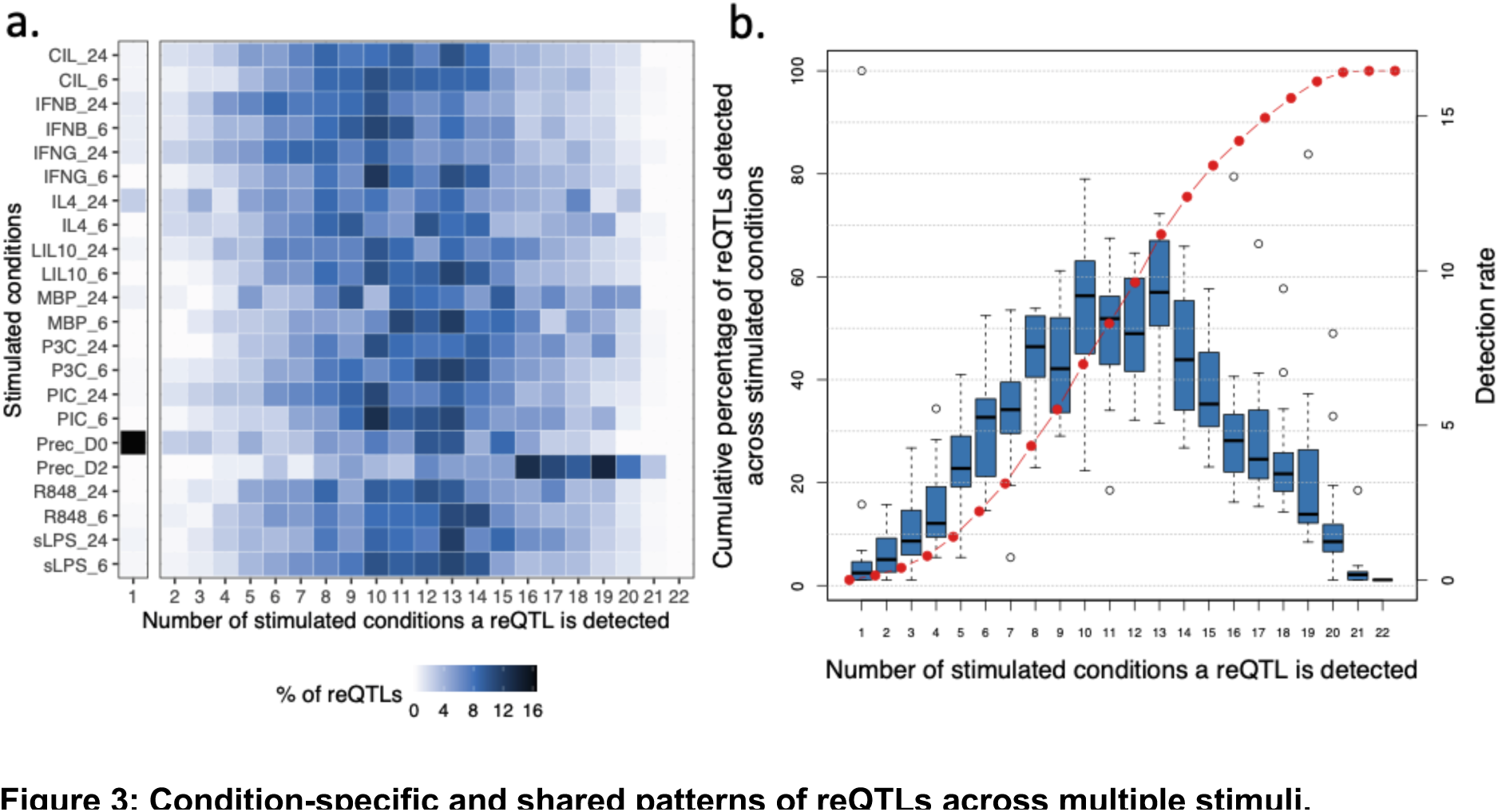
Condition-specific and shared patterns of reQTLs across multiple stimuli. **a.** Proportions of reQTLs for every stimulated condition that were found in a single condition only (column 1) or detected in 2 or more stimulated conditions (columns 2-22). Colour intensity reflects the percentage of reQTLs in a given condition that were also detected in at least one other stimulated condition. Columns represent the number of conditions in which an reQTL was detected **b.** Cumulative percentage of detected reQTLs (red line and points, left hand y axis) and reQTL detection rate (blue boxplots, right hand y axis) versus the number of conditions in which an reQTL was detected.

To investigate the stability of reQTLs over time, we conducted pairwise comparisons between the 6-hour and 24-hour time points within each cell condition, as well as between day 0 and day 2 for precursor cells. To define time point specificity, we examined whether the reQTLs identified in the initial condition (6h,lfsr <=0.05) were absent in the second condition (24h,lfsr >0.05) and vice versa. We observed substantial variation in time-specificity across conditions. For example, cells stimulated with PIC or IFNβ had relatively few time point-specific reQTLs (18-23.4%), perhaps reflecting permanent changes in chromatin state produced by IFNβ stimulation ^30^. In contrast 85% of reQTLs detected in the precursor cells were specific to day, likely reflecting the very substantial changes in transcriptional regulation that occur during cell differentiation (Supplementary Figure 5c).

The GTEx consortium has completed the most extensive and widely used study of eQTLs to date ^3^. The vast majority of GTEx samples were post-mortem tissues and it is unclear how suitable these are for detecting condition-specific genetic effects in immune response. To investigate this, we tested whether the reQTLs detected in our study were also found in GTEx, estimating the overlap using the π1 statistic ^31^ (Supplementary Figure 6). On average, half (54%) of the reQTLs detected in a given condition are replicated in a GTEx tissue, significantly lower than the replication rate of non-response eQTLs (63%) (Supplementary Figure 6c,d) (Wilcoxon P=4.9 x 10^-47^), with the highest replication rates observed for GTEx whole blood (Supplementary Figure 6b) . This suggests that many of the immune pathways we have activated in our study are, at some level, also active in post-mortem tissue.

### Condition-specific genetic variation in disease

We next investigated the relevance of condition-specific eQTLs in disease. We collated a set of 83 well powered (ten or more genome-wide significant loci (P <= 5.0 × 10^-8^) GWAS including 22 immune-mediated, 13 blood-related, 3 cancer, 11 cardiovascular, 15 neurological and 19 other traits or diseases (Supplementary table 9). We found evidence of colocalization ^32^ between a disease association signal and an eQTL for 1955 unique eGenes across all traits and conditions (posterior probability of sharing a single causal variant PP4 >0.75). Relative to the naive conditions, stimulation often increased the strength of evidence of colocalization with disease. We found that including stimulated states substantially boosted our discovery of disease-eQTL colocalisations with only 32% of all colocalisations (631 eGenes) detected in naive cells (Figure 4a).

**Figure 4:**
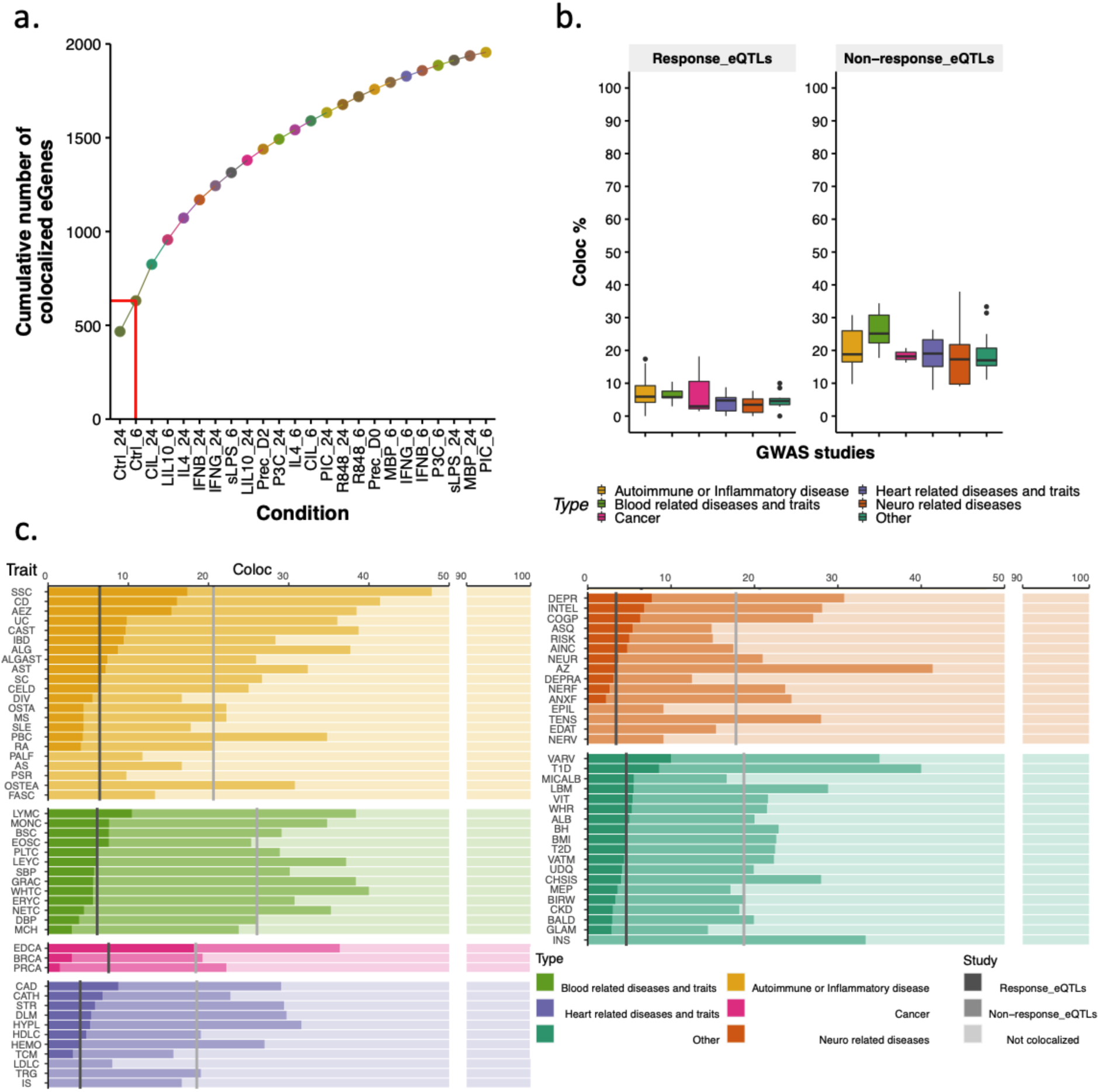
Colocalization of GWAS loci with response and non-response eQTLs across multiple conditions and traits. **a.** Cumulative number of colocalized eGenes seeded in naïve conditions (Ctrl_24,Ctrl_6), with stimulated conditions shown in increasing order. The red line represents the cumulative count of colocalized eGenes exclusively in naive conditions. **b.** Boxplots of percentages of GWAS loci colocalized with response eQTLs and non-response eQTLs across 6 main GWAS categories. **c.** Percentages of GWAS loci colocalized per trait and category. Grey lines indicate the mean percentage of colocalization with response eQTLs across all traits for a specific GWAS category while the light grey line indicates the mean percentage of colocalization with non-response eQTLs.

The rise in the number of eGenes with colocalization evidence can be attributed to both the inclusion of more conditions relevant to disease and the increased power due to the substantial level of biological replication in our study. To investigate this further, we examined the frequency of disease-colocalized eQTLs that had significantly different effect sizes in naive and stimulated conditions using mashr. We found that 21.7% (424/1955; Figure 4b,c) of eGenes with colocalization evidence had at least one reQTL, and among these, 89.4%(379) had a larger effect size in the stimulated condition than in the corresponding naive condition. Disease-colocalizing eGenes showed a significant overrepresentation of reQTL genes (p=0.05, Fisher’s exact test). This suggests that a substantial number of disease-related eQTLs will only be detected in stimulated conditions, because their effect sizes in naive cells are too small.

Our analysis revealed specific traits, as depicted in (Figure 4c), where the use of stimulated macrophages appeared to be more relevant compared to naive conditions. Furthermore, it is important to note that, despite no individual trait exhibiting a substantial increase in the number of eGenes with evidence of colocalization due to reQTLs, this underscores the significance of reQTLs in unravelling disease loci that cannot be explained by naive eQTLs even though their transformative impact on significantly reshaping our comprehension remains limited.

Next,to determine the effectiveness of MacroMap in identifying effector genes within GWAS loci, we compared our results to those obtained using GTEx data. We asked how many of the 1955 disease colocalizations we found in our study could not have been identified using GTEx eQTLs. We found that 998 eGenes (51% 988/1955) were colocalized with higher confidence in our dataset (PP4 > 0.75 in MacroMap versus PP4 < 0.5 across all GTEx tissues) (Supplementary Figure 7), 164 (16.6% 164/988) of which were defined by mashr as reQTLs. All 164 genes were expressed (minimum TPM=0.8, mean TPM across all tissues=165) in at least one GTEx tissue, and had a detectable eQTL (qvalue <=0.05) in one or more GTEx tissues.

### A response eQTL implicates increased expression of *CTSA* with increased risk of coronary artery disease

One such colocalization event was identified between the CAD-associated risk locus at 20q13.12 ^33, 34^ and a reQTL observed 24 hours after stimulation with either sLPS or sLPS+IL10 (LIL10) (PP4=0.98 and PP4=0.99 respectively, Supplementary Figure 8a,d) and not with Ctrl conditions (Supplementary Figure 8b). The risk-increasing allele (rs3827066, C>T) was associated with increased expression of *CTSA* without the gene being differentially expressed after stimulation with either sLPS or LIL10 (Supplementary Figure 8c). Moreover, this risk allele has also been associated with increased risk of Abdominal Aortic Aneurysm (AAA) ^35^, a common complication of vessel wall impairment caused by various predisposing factors including atherosclerosis. Despite these disease associations being known for several years, the disease effector gene in the region had remained elusive. For example, the Open Targets Genetics Portal (https://genetics.opentargets.org/) showed that there are 34 genes in the locus, none of which had a variant-to-gene score >0.3, with *CTSA* ranked as the seventh most likely effector gene in the region. The TSS of *CTSA* is located 67.24kb upstream of rs3827066, with six other genes in closer proximity to the SNP. Data from a promoter capture Hi-C study of circulating immune cells demonstrated that rs3827066 lies in a distal regulatory element of *CTSA* or *NEURL2* in many immune cell types, including monocytes and macrophages ^36^. Our reQTL and colocalization results suggest the disease associated variant acts via dysregulation of *CTSA* expression.

*CTSA* is a lysosomal protective protein with both intracellular and extracellular functions ^37^. In the lysosome, it is essential for both the activity and protection of beta-galactosidase and neuraminidase (*NEU1*) (complex with *CTSA*), an enzyme which cleaves terminal sialic acid residues from substrates such as glycoproteins and glycolipids ^38^. The removal of sialic acid from TLR4, a 2,3 sialylated pattern recognition receptor, with the action of *NEU1* ^39, 40^, facilitates LPS recognition and the subsequent downstream activation of NF-κB signalling and inflammatory cytokine production. Outside the cell, *CTSA* has been suggested to play a role in extracellular matrix (ECM) remodelling ^41–44^. ECM remodelling is an integral part of several chronic inflammatory processes including atheromatous plaque formation within vessel walls, and subsequent CAD ^45^. The cardiac expression of *CTSA* is upregulated in multiple animal models of myocardial infarction, Type 2 Diabetes and angiotensin II-stimulated hypertrophy ^46–49^. Increased expression of *CTSA* has been shown to trigger proteolysis of the extracellular antioxidant enzyme EC-SOD, resulting in higher levels of oxidative stress, myocyte hypertrophy, ECM remodelling, and inflammation ^44, 50^. We thus hypothesise that rs3827066 increases risk of CAD and AAA by disrupting a *CTSA* regulatory element, leading to increased expression of *CTSA* and elevated oxidative stress during the ECM remodelling stage of atheromatous plaque formation. Detecting this *CTSA* eQTL only upon stimulation (i.e. reQTL) is consistent with ECM remodelling being induced following a prolonged period of vessel wall inflammation in CAD. Overall, our findings suggest that this CAD-associated risk locus exacerbates oxidative stress in the ECM by abnormally upregulating CTSA following an inflammatory response.

## Discussion

In this study, we used a high-throughput cellular system of human IPSC-derived immune cells to survey inter-individual variation in gene expression across a range of stimulated conditions. We identified 10,170 eGenes, substantially more than previous eQTL studies with similar sample sizes. The increase in eQTL discovery resulted from the combination of higher depth expression profiling, and enhanced detection of both condition-specific gene expression and the condition-specific effects of genetic variants. Using a statistical method that accounts for low power, we detected a significant change in genetic effect size between naive and stimulated cells in 23% of all eGenes. For this set of response eQTLs we found that the majority were shared widely across stimulated conditions, with effects found in a single condition constituting a small fraction (1%). We discovered 1955 disease-eQTL colocalizations of which 51% were not detectable in any GTEX tissue suggesting many disease-associated variants may function in a condition-specific manner.

The number of eQTLs we detected in this study was higher than might be expected given our sample size. Perhaps surprisingly, our results suggest that a high level of biological replication was at least as important as condition-specific genetic effects in boosting our discovery rate. For example, profiling naive cells at two different timepoints yielded an additional 774 new eQTLs. More generally, it is unclear how many eQTLs could have been detected by a better powered study of naive cells rather than profiling a large number of stimulated conditions. Nonetheless, profiling stimulated cells proved valuable for interpreting disease loci. We found that 21.7% (617) of all disease colocalizations had a different effect size, usually greater, in a stimulated condition compared to naive cells. However, we believe our estimate for the true number of response QTLs is likely to be a lower bound, mainly limited by statistical power. In some cases, the statistical method we used performed aggressive shrinkage of eQTL effect sizes towards zero. For example, we observed colocalization of a *CD80* eQTL after stimulation with sLPS at 6 hours that was not confidently colocalized in naive cells. Despite this discrepancy in colocalization confidence between naive and stimulated macrophages, this eQTL was not considered a reQTL by mashr (Supplementary Figure 9 and 10). We expect that future studies with larger sample sizes will increase the number of eQTL effects that can be confidently considered response eQTLs. Several recent studies suggest that a smaller than expected fraction of GWAS loci colocalize with an eQTL ^3, 5^ either because genetic effects are often restricted to cell types and conditions that are not causal for the trait ^6^. MacroMap represents the most comprehensive characterization of macrophage eQTLs in different environmental contexts to better understand how genetic variants impact human traits. Our findings highlight the value of reQTLs for expanding our understanding of complex diseases. While their transformative impact on significantly reshaping our comprehension remains limited, reQTLs offer unique insights into the intricate genetic regulation that surpass what naive eQTLs can provide. Their contribution to uncovering previously unexplained disease loci is undeniable, paving the way for further exploration and bringing us closer to a comprehensive understanding of the complex genetic mechanisms underlying the disease. Despite MacroMap being able to uncover a significant proportion of “missing” disease-relevant eQTLs, many remain to be discovered. Upcoming large-scale single-cell QTL and response QTL studies of primary tissues and appropriate cell models will further advance our ability to detect disease-relevant genetic effects. Well-powered cell type and context specific QTL studies of molecular traits with different genomic properties (for example, splicing, chromatin accessibility and chromatin interactions) will likely further improve our ability to understand human disease biology.

## Methods

### iPSC lines

Human iPSCs lines from healthy donors of European descent were selected from the HipSci project ^51^ (http://www.hipsci.org) for differentiation to macrophages (see supplementary note). All HipSci samples were collected from consenting research volunteers recruited from the NIHR Cambridge BioResource (http://www.cambridgebioresource.org.uk). Briefly, 315 lines were initially selected and 227 of them (71.6%) were successfully differentiated. RNA-seq libraries were produced for 217 lines and based on quality control 209 unique lines (Supplementary table 10) (4698 unique RNA-seq libraries across all conditions) were included in the final dataset (supplementary Figure 1A,B).

### RNA-seq and quality control

To process the large number of libraries more efficiently, two RNA-seq library construction protocols were utilised, including a modified Smart-seq2 protocol and the NEBnext Ultra II Directional RNA Library kit (further details provided in the supplementary note). However, this resulted in a batch effect due to the different library preparation methods. This effect was included as a covariate in all downstream analyses.

RNA-seq reads (75 bp paired-end) were aligned to the GRCh38 reference human genome and gencode v.27 transcript annotation using STAR_2.5.3a ^52^. To quantify gene expression we used featureCounts v1.5.3 ^53^. We kept protein-coding and lincRNA genes in all analyses with mean expression >= 0.5 transcripts per million (TPMs) in at least half of the conditions (>=12), resulting in a total of 14,060 genes. To ensure the quality of the samples, we employed several QC metrics. Principal Components Analysis (PCA) was performed per 96-library pool (4 iPSC lines per pool, 24 conditions per iPSC line) to detect sequencing outliers. Non-stimulated or mislabeled labelled stimulated samples were identified and discarded based on pairwise PCA comparisons of each condition with the rest of the conditions, per 96-library pool. Sex incompatibility checks were also performed using the methods described in ^54^ and 3 iPSC-lines (72 samples) were discarded due to discordant sex annotations. Subsequently, we performed UMAP analysis ^55^ to cluster the different conditions and wrongly labelled samples that passed PCA filtering were discarded. FInally, we utilised the Match BAM to VCF (MBV) method ^56, 57^ to detect sample swaps and cross contamination between RNA-seq samples. We discarded 3 iPSC-lines (72 samples) and 63 additional samples due to cross contamination, and corrected the labels for 23 iPSC-lines identified as swaps. We did not observe concordance of genotype-RNA-seq data for 4 lines which we kept in the final dataset for differential expression analysis but discarded from eQTL mapping. Among the 23 swaps, 2 lines were identical with lines already present in the data and were subsequently removed from the dataset.

In total, we discarded 8 iPSC-lines (∼3.7% of the successfully differentiated lines) and 510 RNA-seq samples (∼9.8%) based on our QC metrics (318 samples based on all QC metrics, 192 from the discarded 8 iPSC-lines, Supplementary Figure 1C) resulting in a total of 4698 unique RNA-seq libraries across all conditions.

### Variance component analysis

We adopted the same approach that was implemented in ^60^ to quantify transcriptional variation. In brief, we used a linear mixed model that employed log10(TPM +0.1) values for the 14060 genes, with 15 technical confounders (including RunID, Donor, Stimulus Hours, Sex, Library preparation method, Date thawed, Passage number at thawing, Passage EB formation, IPSCs culture time, Total Harvests, Differentiation time No of Days, Purity results, Estimated cell diameter, SD cell diameter, and Differentiation media) fitted as random effects with independent variance parameters 𝜑^2^_*k*_. We measured the variance explained by factor k using the intraclass correlation 𝜑^2^_*k*_/(1+𝜑^2^_*k*_), while the remaining 14 factors were held constant. The standard error of the intraclass correlation was computed using the delta method, with the standard error of the variance parameter estimator.

### Genotypes

GRCh37 imputed genotypes were obtained from the HipSci project ^51^. We utilised CrossMap ^58^ to lift over the variant coordinates from GRCh37 to GRCh38. We then used bcftools to filter the resulting VCF file, retaining only variants with INFO score >0.4 and minor allele frequency (MAF) > 0.05. To address population stratification, we used EIGENSTRAT ^59^ to calculate genotype principal components (PC) for the retained variants.

### UMAP clustering and visualisation

To visualise the transcriptional variation across conditions, we applied UMAP analysis to the gene expression data. Prior to UMAP, we performed several preprocessing steps on the log-transformed transcripts per million (TPMs). First, we quantile-normalised the log-TPMs to remove technical differences between samples. Next, we applied a rank-based inverse normal transformation to ensure that the gene expression values were normally distributed. Finally, we regressed out (linear regression) the effects of several covariates including runID, donor, library preparation method, sex, purity results, differentiation media, estimated cell diameter, and Differentiation time No Days (Time in days from EB plating until the day of successful harvest) to account for technical variation and batch effects. The resulting UMAP plot provided a low-dimensional visualisation of the transcriptional differences among the different conditions.

### Differential gene expression analysis

DESeq2 ^22^ was used to identify differentially expressed genes between the naive and stimulated conditions, and SVA (surrogate variable analysis) ^61^ was employed to detect hidden technical variation that could not be captured by our technical covariates. Specifically, we fitted the samples from both the 6 and 24 hour time points of the stimulated and naive conditions together and included 10 SVA factors that were determined from the overall sample composition. An interaction term was also included in the model as shown below:

DESeqDataSet(group1_vs_group2,design= ∼ X1 + X2 + X3 + X4 + X5 + X6 + X7 + X8 + X9 + X10 + Stimulus + Hours + Stimulus:Hours).

Following this, the same model was fitted without the interaction term and a likelihood ratio test (test=“LRT”) was performed to compare the full model (including all SVA factors and the interaction term) to the reduced model (including all SVA factors but not the interaction term).

To identify differentially expressed genes at specific timepoints (either 6 or 24 hours) or genes showing different differential expression patterns between time points (interaction term, time point effects), we used the Wald test (test=“Wald”, alpha=0.05) in DESeq2. Finally, we assessed significance at a 5% false discovery rate (FDR) using the Benjamini-Hochberg and kept genes with abs(log2FoldChange) >= 1.

### eQTL mapping

We mapped cis-eQTLs within ±1Mbp of the transcription start site (TSS) of each gene using QTLtools v.1.1 ^57^. Briefly, QTLtools conducts permutations of the expression data for each gene to record the best P value for any SNP in the cis window. The distribution of the best P values follows a beta distribution under the null hypothesis, and QTLtools estimates the parameters of the beta distribution of each gene through maximum likelihood, which depends on the LD structure of the cis region. An adjusted gene-level P value is computed based on the beta distribution for each gene. To correct for multiple testing across all genes, we used the q-value R package ^62^ on the adjusted gene-level P values obtained from 1000 permutations and significance was assessed at 5% FDR (qvalue <0.05) to identify genes with at least one significant cis-eQTL (“eGenes”). We included expression PCs (35-50 depended on condition) and 3 genotyping PCs as covariates to correct for technical variation and capture population stratification.To determine the optimal configuration (number of expression PCs per condition) that maximised the number of discoveries (eGenes), we repeated the entire analysis multiple times using different numbers of expression PCs. Multiple independent signals (5% FDR) for a given eGene were identified by forward stepwise regression followed by a backwards selection step implemented in QTLtools (conditional pass).

### Functional enrichment analysis

We performed an enrichment analysis of genomic annotations to investigate the functional implications of our identified eQTLs. Firstly, we utilised Ensembl’s Variant Effect Predictor (VEP) and the Ensembl Regulatory Build to annotate the eQTLs. To identify specific genomic annotations enriched among our eQTLs, we used the first stage hierarchical model implemented in PHM ^63^ (https://github.com/natsuhiko/PHM).

### Response eQTLs using Multivariate Adaptive Shrinkage (mash)

In order to determine which eQTLs in our dataset were truly restricted to stimulation (reQTLs) we used mashr ^28^ by following the workflow provided by the authors of mashr(https://stephenslab.github.io/mashr/articles/eQTL_outline.html). Initially, we calculated the standard errors of QTL effect sizes (betas) from QTLtools nominal output, which were combined with effect sizes as input data for mash. Our analysis consisted of two subsets of tests: a random subset of 200,000 tests comprising both null and non-null tests, and a more focused “strong” subset that specifically included the lead SNP (lowest p-value) per gene across all conditions, emphasising the most impactful associations.While the strong subset of tests was used to learn data-driven covariance matrices, the random subset of tests was used to estimate mixture weights and scaling coefficients, as well as to learn the correlation structure among null tests. Additionally, we employed the mashr mode “mashr with common baseline,” as described here (https://stephenslab.github.io/mashr/articles/intro_mashcommonbaseline.html), by setting Ctrl_24 as our baseline condition and excluding Ctrl_6 from the mash analysis. In the common baseline mode, mashr estimated the deviation of eQTL effect sizes in each alternative condition from that of the baseline condition, taking into account the correlation that arises when comparing all conditions to a common baseline. To execute the analysis, we first fitted the mashr model to the random tests to determine the mixture weights. We then utilised the model fit to compute the posterior mean effect sizes (mash effect sizes) on the best associated SNP per gene for every stimulation condition. We considered significant response eQTLs (reQTLs) gene-SNP pairs with a local false sign rate (LFSR) below 0.05. The LFSR is a measure that is stricter than the false discovery rate (FDR) since it not only requires significant discoveries to have a nonzero value, but also to have a consistent sign ^29^. To determine the reQTLs that have consistent effects across multiple conditions (shared reQTLs) or function in only a single condition (condition-specific reQTLs), we needed to investigate whether the gene-SNP pair for a particular condition had LFSR <0.05 in the other conditions. This allowed us to quantify the level of sharing of response effects among the different conditions. For instance, if a particular gene-SNP pair was significant in three out of four stimulation conditions, we considered it a shared reQTL across those three conditions. Conversely, if a gene-SNP pair was significant only in one condition, we classified it as a condition-specific reQTL for that particular condition.

### Colocalization

Colocalization analysis was performed with coloc v3.2-1 ^32^ between our eQTL summary statistics and 83 publicly available GWAS summary statistics (either from GWAS catalogue or from UK BioBank GWAS ^64^). These GWAS represented 22 immune-mediated, 13 blood-related, 3 cancer, 11 cardiovascular, 15 neurological, and 19 other traits or diseases (Supplementary table 9). To ensure that we only included datasets with sufficient statistical power, we only considered GWAS datasets that had ten or more genome-wide significant regions (P <= 5.0 × 10^-8^). Specifically, to identify significant regions, we created bed files for all SNPs that had (P <= 5.0 × 10^-8^) for each GWAS study. We then used the ’bedtools merge -d 500000’ command to combine overlapping variants into a single region that spanned all the combined variants. The regions were expanded by 500 kb on either side, and any overlapping regions were merged again. Next, we run coloc on a 2 Mb window centred on each lead eQTL, using default priors and set the colocalization threshold as PP4 > 0.75. The colocalization proportions were calculated as the proportion of colocalized significant regions among all identified significant regions per GWAS.

## Acknowledgements

We would like to thank the Human Genetics Informatics team at the Wellcome Sanger Institute for providing computational support to run the analyses described in this manuscript. This work was supported by Wellcome Sanger Institute Core funding from the Wellcome Trust (206194, 220540/Z/20/A). The iPSC lines were generated at the Wellcome Sanger Institute, under the Human Induced Pluripotent Stem Cell Initiative funded by a strategic award (WT098503) from the Wellcome Trust and Medical Research Council. NIP was supported by the Early Postdoc Mobility fellowship from the Swiss National Science Foundation.

## Author Contributions

NIP performed the analyses and, along with OEG, CAA and DJG drafted the manuscript with contributions from all authors. OEG assisted with multiple analyses, independently validated eQTL results and provided feedback to improve the manuscript. JR performed the conditional eQTL analysis. NK provided statistical feedback on several occasions, assisted with multiple analyses, and conducted analysis on the pilot data. MI, LBV, AT and CG applied the differentiation protocol, performed QC metrics on the differentiated macrophages and carried out the stimulations under the supervision of CG. AK optimised the low-input bulk RNA-seq protocol and prepared the RNA-seq libraries along with MI and AB. DJG conceptualised the study while CAA and DJG supervised this work.

## Data availability

Imputed genotype data for the HipSci lines are available from ENA (xxxx) and EGA(xxxxx). Unprocessed RNA-seq data are available from ENA (xxxxx) and EGA (xxxxx). Full summary eQTL statistics are available from Zenodo (https://doi.org/10.5281/zenodo.7967759).

## Supplementary note

### iPSC culture and macrophage differentiation

iPSC culture and macrophage differentiation was carried as previously described ^15^ but with some minor modifications: embryoid bodies were harvested 3 days after formation and transferred onto gelatinised tissue-culture treated 10 cm dished in serum-free X-VIVO 15 (Lonza, BE02-060F) or Stem Pro-34 SFM (Thermo Fisher, 10640-019), with both mediums supplemented with 2 mM GlutaMAX (Thermo Fisher, 35050061), 50 IU/ml penicillin, 50 IU/ml streptomycin (Sigma, P4333), 100 ng/ml human macrophage colony stimulating factor (hM-CSF) (Peprotech, 300-25) and 25 ng/ml human interleukin-3 (hIL-3) (Peprotech, 200-03). Macrophage progenitor cells were counted and plated in macrophage complete media (RPMI 1640 (Thermo Fisher, 11875093) supplemented with 10% heat-inactivated FBS (Thermo Fisher, 10500-064), 2mM GlutaMAX (Thermo Fisher, 35050061) and 100 ng/ml hM-CSF (Peprotech, 300-25)) at a cell density of 10,000 cells per well on a 96-well plate (for RNA-seq), or 25,000 cells per well of black 96-well plate (VWR, 734-1661) (for the macrophage purity assay) and differentiated for another 7 days.

### iPSC-derived macrophage purity assay

iPSC-derived macrophages progenitor cells were seeded and differentiated as above before fixing in 50 µL of 4 % formaldehyde (Applichem, A0823.2500) at 4°C for 20 minutes. Cells were washed twice in 100 µL PBS with calcium and magnesium (Sigma, D8662) before blocking in 10% (v/v) donkey serum (AbD Serotec, C06SBZ) 0.1% Triton X-100 (Sigma, 93420) at room temperature for 1 hour. Cells were then stained with 1:200 anti-CD14 (BioLegend, 301802) and 1:800 anti-CD68 (Cell Signalling Technology 76437S) in 1% blocking solution overnight at 4C. We then washed the cells three times with PBS and stained with secondary antibodies (1:1000 donkey anti-mouse AF647 and 1:1000 donkey anti-rabbit AF488) and DAPI (10ug/mL, AppliChem A1001) at room temperature for 1 hour. Wells without primary antibody were used as negative staining controls. Cells were washed three times and imaged on a Cellomics Arrayscan (ThermoFisher), and the proportion of CD14+CD68+ cells calculated. Only cell lines with greater than 90% double stained cells were processed for RNAseq.

### iPSC-derived macrophage stimulation conditions

After the 7-day differentiation to macrophages, cells were incubated for 6 or 24 hours in complete macrophage media on its own (controls) or containing the following stimuli: : 10 ng/mL recombinant human interleukin-10 (Peprotech, 200-10-2), 10 ng/mL recombinant human interferon-b (Peprotech, 300-02BC-5), 20 ng/mL recombinant human interleukin-4 (Peprotech, 200-04-5), 50 ng/mL P3C (Pam3CSK4) (Tocris, 4633/1), 20 ng/mL recombinant human interferon-γ (Peprotech, 300-02-20), 10 ng/mL lipopolysaccharides from Escherichia coli O127:B8 (Sigma Aldrich, L3129), 40 ng/mL human recombinant tumour necrosis factor alpha (Peprotech, 300-01A-10), 100 ng/mL R848 (Resiquimod) (Invivogen, tlrl-r848), or 5 ng/mL recombinant human sCD40 Ligand (Peprotech, 310-02-10). For the stimulations with HMW poly I:C (Invivogen, tlrl-pic), macrophages were transfected with poly I:C as follows: 0.15 mL of 1 mg/mL of poly I:C was mixed with 0.3 mL P3000 reagent (Lipofectamine 3000 kit) and 5 mL Opti-MEM (Thermo Fisher, 31985062). In another tube, 0.3 mL of Lipofectamine 3000 (Thermo Fisher, L3000001) and 5 mL of Opti-MEM were mixed. The diluted Lipofectamine 3000 and poly I:C tubes were mixed and incubated at room temperature for 10 minutes to allow complexes to form. 100 mL of macrophage differentiation media from above was added to the poly I:C complexes and mixed. Media was removed from the macrophages and replaced with the diluted poly I:C complexes. Control transfections were carried out in exactly the same way but without the addition of poly I:C.

### iPSC-derived macrophage low-input bulk RNA-seq preparation

At the end of the macrophage stimulation period, the media was removed and cells were lysed immediately by adding 50 µL of a 1x lysis/binding buffer (100 mM Tris-HCl pH 7.5, 0.5 M LiCl, 10 mM EDTA, 1 % w/v lithium dodecyl sulphate, and 5 mM 1,4-dithiothreitol) and mixed well. Lysed cells were stored at -80 °C until needed. Using the automated Zephyr G3 NGS Workstation (Perkin Elmer), mRNA was purified from the cell lysates in 96 well plates using the mRNA DIRECT kit (Thermo Fisher, 61012), according to the manufacturer’s instructions, using 20 µL of oligo dT Dynabeads. The purified mRNA was eluted in either 7 µL of nuclease-free 10 mM Tris-HCl pH 7.5 for processing through the modified Smart-seq2 method, or 5 mL of nuclease-free water for processing through the NEBnext Ultra II Directional RNA Library kit (E7760L). For the modified Smart-seq2 method ^65^), the purified mRNA was processed as follows : 2 µL of oligo dT_30_VN (Integrated DNA Technologies) and 2.34 µL of 10 mM dNTPs (Thermo Fisher, R0193) were mixed with 7 µL of the purified mRNA and heated to 72 °C for 3 minutes to denature secondary structures, before rapidly chilling on ice for 5 minutes. 5 µL of 5x SMARTScribe first-strand buffer (Clontech Takara, 639538), 0.63 µL of SUPERase inhibitor (Thermo Fisher, AM2696), 1.25 µL of 100 mM 1,4-dithiothreitol, 5 µL of betaine (Sigma, B0300-5VL), 0.15 µL of 1 M MgCl_2_, 0.38 µL of template-switching LNA-oligo (TSO) (Qiagen) and 1.25 µL of SMARTScribe reverse transcriptase (Clontech Takara, 639538) were added to the denatured mRNA/dNTP/oligo dT_30_VN mix. Following a brief vortex mix, reverse transcription was performed at 42 °C for 90 minutes, followed by 10 cycles of 50 °C for 2 minutes, then 42 °C for 2 minutes. The reaction was stopped by incubating at 70 °C for 15 minutes. The first-strand cDNA was purified using 0.8 volumes of Ampure XP beads (Beckman Coulter, BCAG0006) to 1 volume of the reverse transcription reaction volume, according to the manufacturer’s instructions, but leaving the eluted cDNA in 12 µL of 10 mM Tris-HCl pH7.5 with the beads in solution. This was done to maximise the amount of cDNA carried forward to the subsequent cDNA amplification reaction. The cDNA was amplified by adding 0.5 µL of 10 µM ISPCR primer (Integrated DNA Technologies) and 12.5 µL of 2x KAPA HiFi polymerase (Kapa Biosystems, KK2601) to the 12 µL of cDNA and mixed before heating at 98 °C for 3 minutes, followed by 11 cycles of 98 °C for 20 seconds, 67 °C for 15 seconds and 72 °C for 6 minutes, followed by a final extension at 72 °C 5 minutes. The amplified double-stranded cDNA was purified as before, but this time the Ampure XP beads were removed from the 20 µL eluate. Amplified double-stranded cDNA was quantified with a Quant-iT^TM^ dsDNA high sensitivity assay kit (Thermo Fisher, Q33120) in black v-bottom 96-well plates (Greiner Bio-One, 651209) on a FLUOstar Omega (BMG Labtech), according manufacturers’ instructions. For cDNA tagmentation, 4 ng of cDNA was diluted with 10 mM Tris-HCl pH 7.5 to a volume of 9.5 µL. 5 µL of a 3x tagmentation buffer (99 mM Tris acetate, 198 mM potassium acetate, 30 mM magnesium acetate and 48 % v/v N,N-dimethylformamide) and 0.5 µL of TDE1 (Illumina, 20034197) were added, mixed and incubated at 55 °C for 5 minutes. The tagmentation reaction was stopped by the addition of 2.5 µL of a tagmentation stop buffer (220 mM EDTA and 1.1 % w/v sodium dodecyl sulphate) and mixed before incubating at room temperature for 10 minutes. The tagmented cDNA was diluted with 10 mM Tris-HCl pH 7.5 to a final volume of 50 µL, before purifying with a 2:1 ratio of Ampure XP beads to sample volume, eluting the tagmented cDNA in 7 µL of 10 mM Tris-HCl pH 7.5. Tagmented cDNA samples were then amplified and sample-indexed by PCR as follows: 7 µL of tagmented cDNA was added to 2.5 µL of i5 index adapter and 2.5 µL of i7 index adapter from the Nextera® XT index kit v2 set A (Illumina, 15052163), 0.25 µL of 50 µM PC1 primer, 0.25 µL of 50 µM PC2 primer and 12.5 µL of 2x KAPA HiFi polymerase, before mixing and incubating at 72 °C for 3 minutes, 98 °C for 30 seconds, followed by 9 cycles at 98 °C for 15 seconds, 62 °C for 30 seconds and 72 °C for 30 seconds, followed by a final extension at 72 °C for 3 minutes. Individual libraries were purified, excess primers removed by performing 0.8:1 ratio of Ampure XP beads to PCR volume, eluting the finished library in 20 µL of 10 mM Tris-HCl pH 7.5.mRNA processed through the NEBnext Ultra II Directional library kit was done so according to the manufacturer’s instructions with 17 cycles of PCR.

All libraries were quantified with a Quant-iT^TM^ dsDNA high sensitivity assay kit, as mentioned above, before combining 96 libraries per pool in equimolar amounts. Library pools were assessed for fragment length and quantity on a Bioanalyser using a High Sensitivity DNA kit (Agilent Technologies, 5067-4626), according to the manufacturer’s instructions. Each 96-library pool was sequenced over 8 lanes of a HiSeq SBS v4, collecting 75 bp paired-end reads

## Supplementary figures

**Supplementary figure 1:**
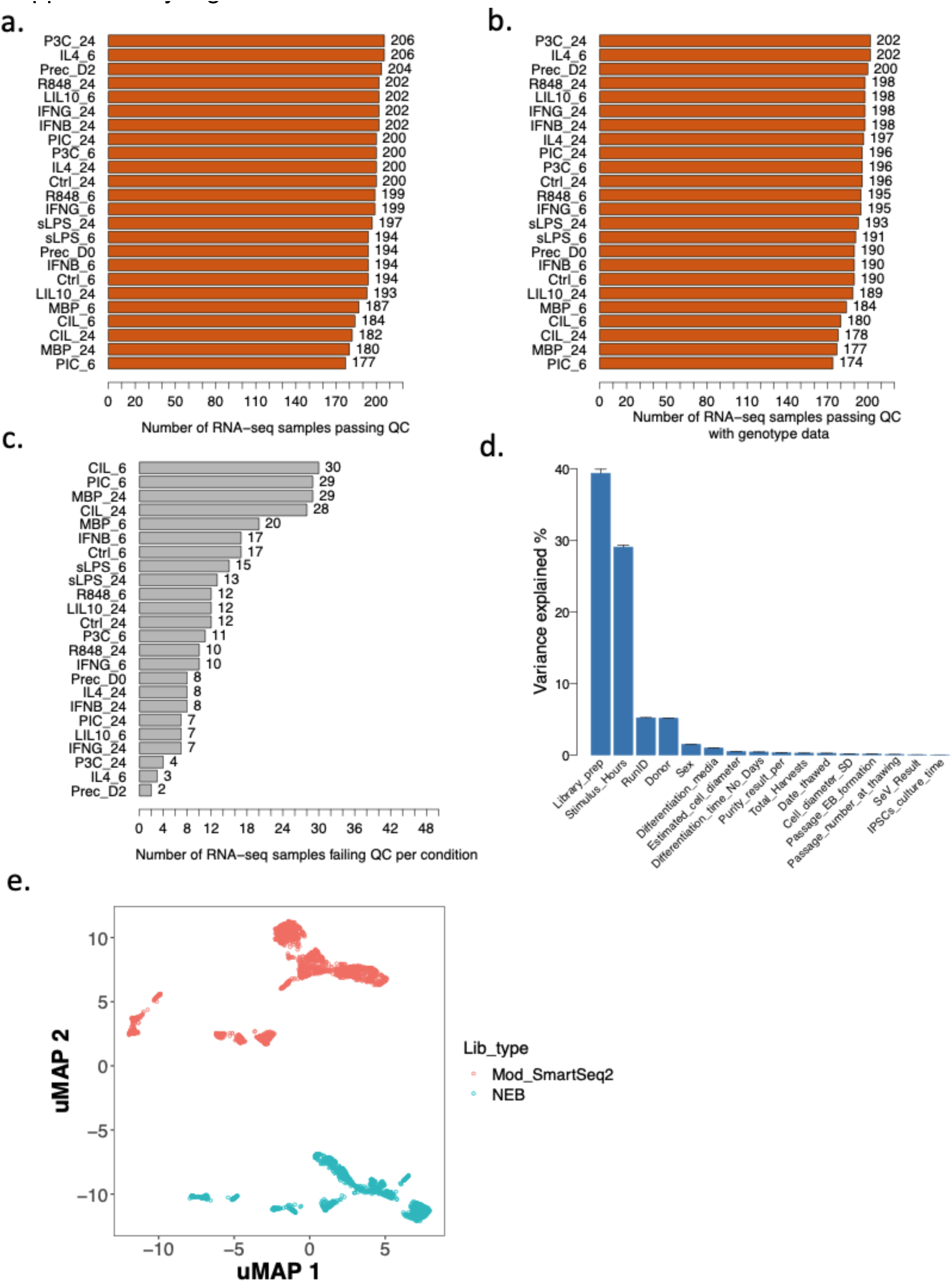
Comprehensive assessment of RNA-seq data quality. **a**. Number of RNA-seq samples per condition that passed the quality control metric (Methods). **b.** Number of RNA-seq samples that had matching genotype data per condition. **c.** Number of RNA-seq samples per condition failing quality control metrics (Methods) **d.** Variance deconvolution analysis taking into account multiple technical and biological covariates that could influence gene expression. **e.** UMAP representation of gene expression data without regressing out the Library Preparation method. The use of two different library preparation protocols, modified SmartSeq2 and NEBnext Ultra II Directional RNA Library kit (NEB), caused a strong batch effect that is clearly captured in the UMAP 2 and is included as a covariate in all downstream analyses.

**Supplementary figure 2:**
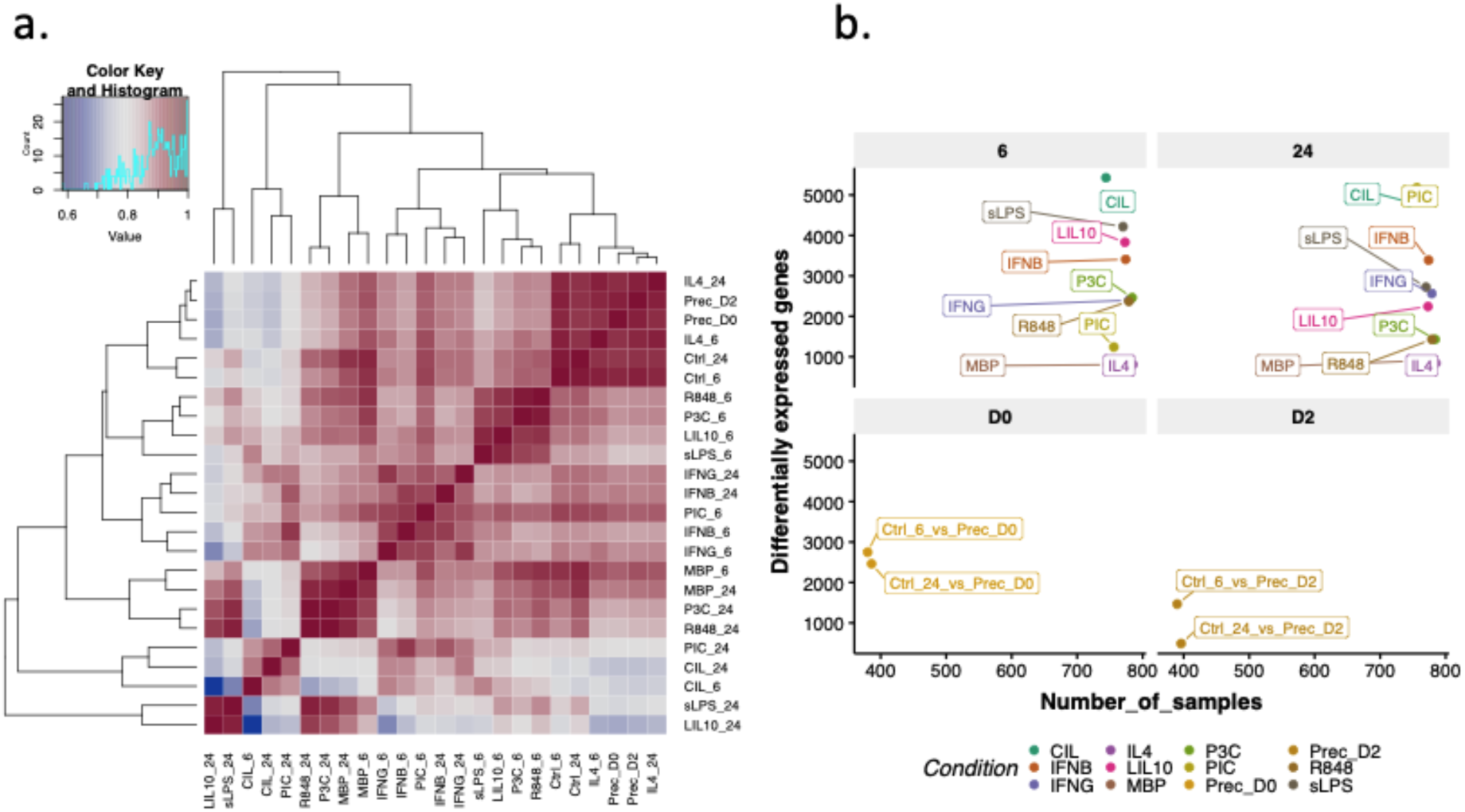
Impact of stimulation on gene expression: differential expression analysis and gene expression correlation clustering. **a**. Heatmap and clustering of correlation of expression of quantified genes (mean TPM values per gene across all individuals) for a given condition. **b.** Number of differentially expressed genes per condition after 6 and 24 hours of stimulation compared to naïve conditions. The differentially expressed genes were defined at false discovery rate (FDR) = 5% and fold change ≥2.

**Supplementary figure 3:**
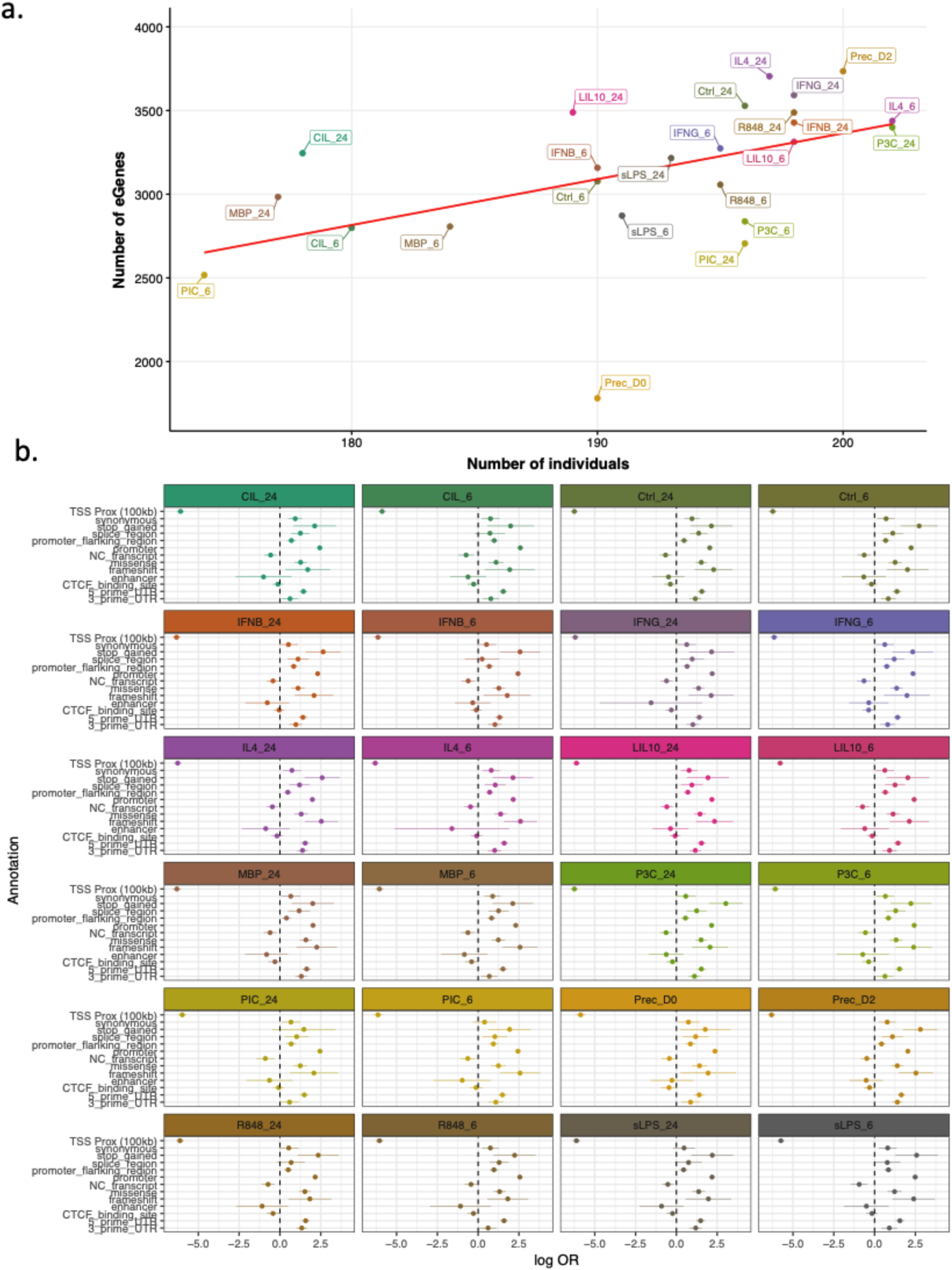
eQTL discovery and its functional implications. **a.** Number of genes with a cis-eQTL (eGenes) per condition as a function of the sample size. **b.** Enrichment analysis of eSNPs for each condition in functional annotations.Enrichment is shown as logOR. The depletion of TSS proximity means that every 100Kb further away from the TSS the number of eSNPs is decreasing.

**Supplementary figure 4:**
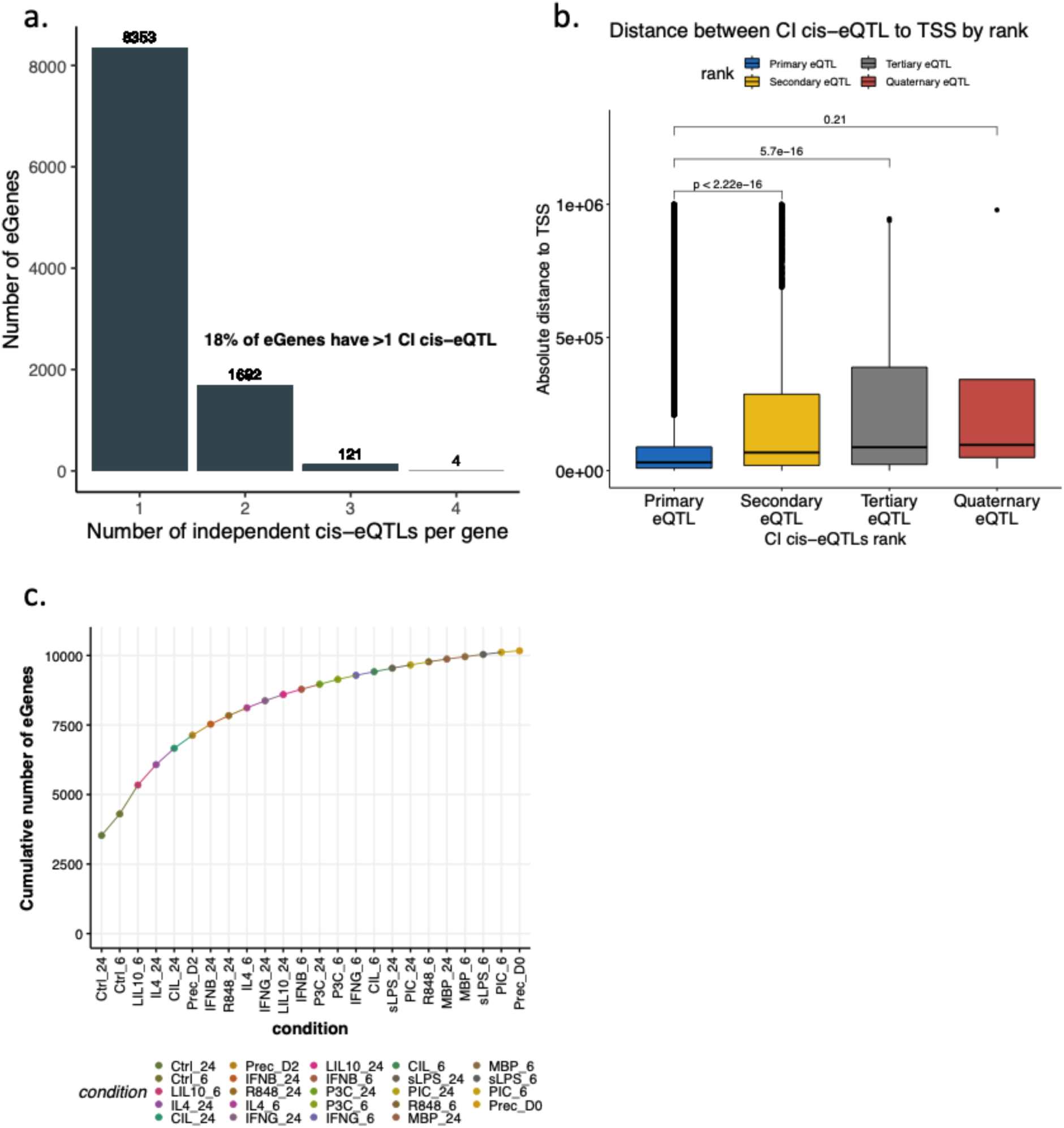
Total number of eGenes across conditions and conditional eQTL mapping. a. Number of eGenes with multiple independent cis-eQTLs based on conditional analysis (Methods). b. Distribution of distances of conditionally independent eQTLs from the transcription start site (TSS) of their corresponding eGenes. c. Cumulative number of eGenes seeded in naïve conditions (Ctrl_24,Ctrl_6), with stimulated conditions shown in increasing order.

**Supplementary figure 5:**
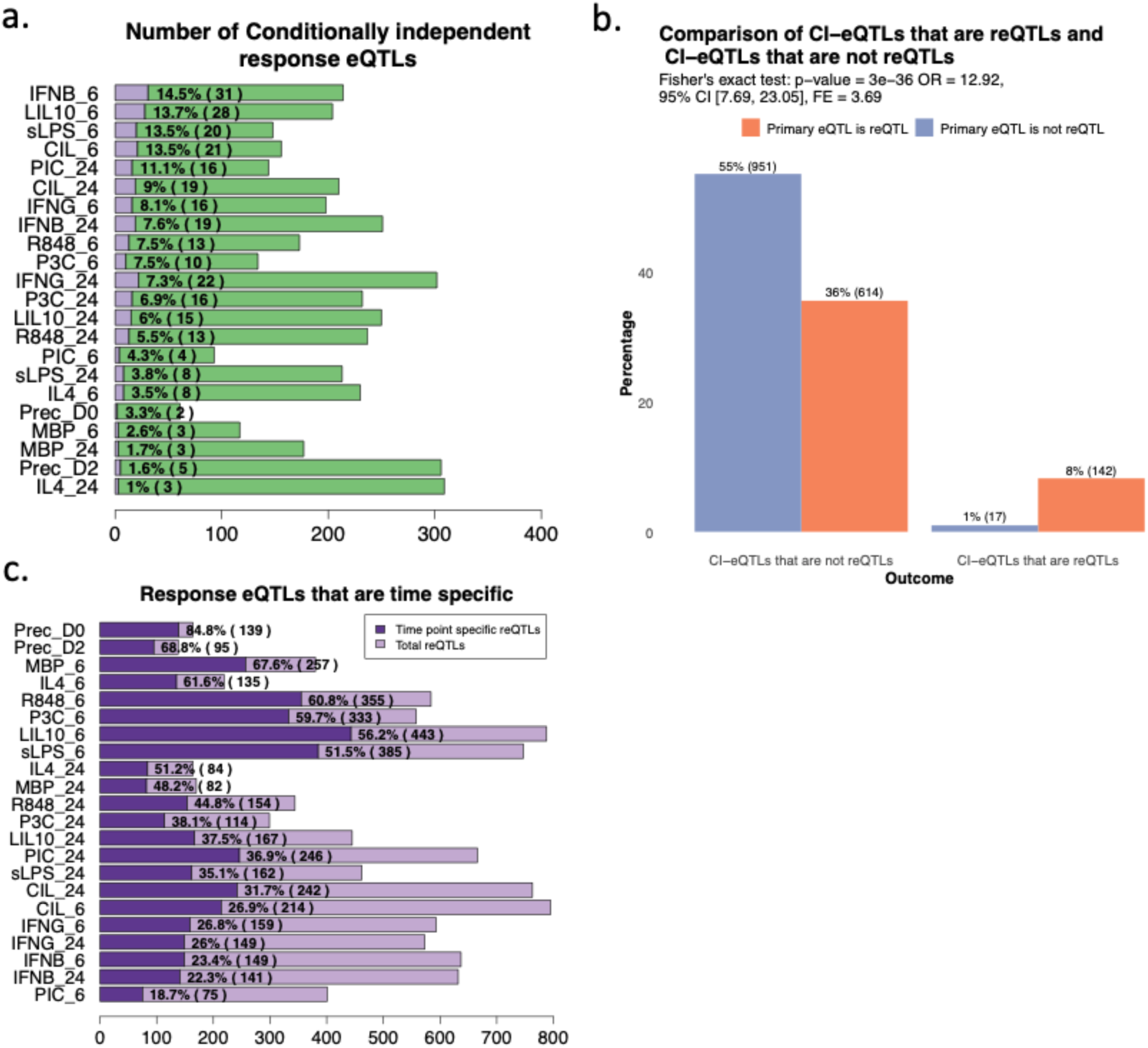
Landscape of conditionally Independent reQTLs and reQTLs across different time points. **a.** Number and proportions of conditionally independent reQTLs (purple) per stimulation condition ordered from highest to lowest proportion. **b.** Enrichment of conditionally independent eQTLs (CI-eQTLs) that are reQTLs compared to those that are not reQTLs. Out of a total of 159 genes with a conditionally independent CI-reQTL, 142 genes (89%) also had a primary reQTL. This represents an almost 4-fold enrichment, with Fisher’s exact test p-value =3^-36^ and an odds ratio of 12.92 (95% confidence interval 7.7 -23.05). **c.** Number and proportions of time point specific response QTLs ordered from highest to lowest proportion.

**Supplementary figure 6.**
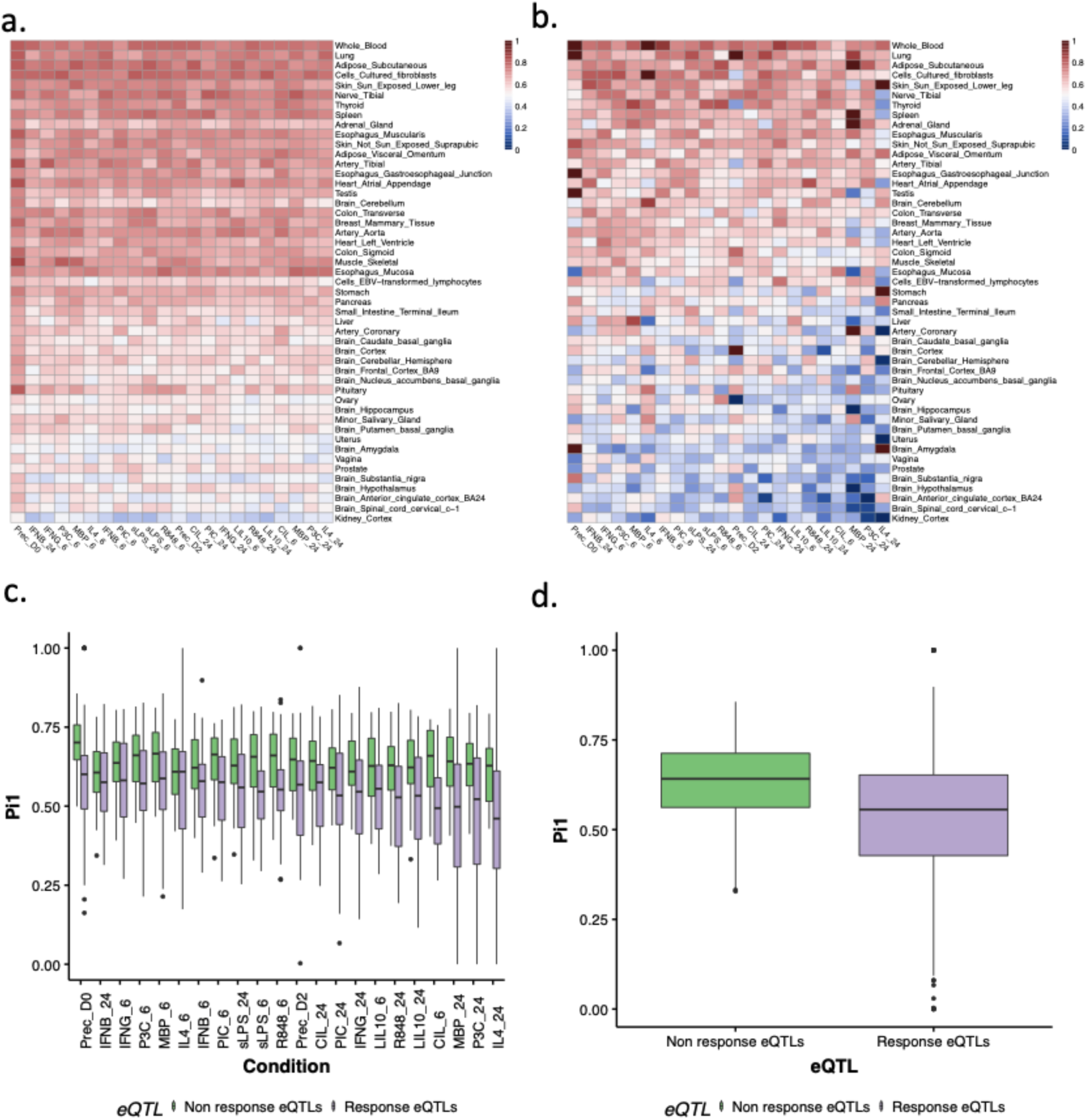
Detailed information on the replication rate of non-response and response QTLs-(reQTLs). **a.** Heatmap displaying π1 values for non response eQTLs and GTEx tissues. The π1 values are arranged in order from tissues with the highest replication rate to those with the lowest. Higher π1 values indicate a higher replication rate. **b.** Ηeatmap following the same ordering as in panel (a), but for response eQTLs in all GTEx tissues.**c.** Distribution of π1 values for non response (green) and reQTLs (purple) per condition. **d.**. Distribution of the mean π1 values (across all GTEx tissues) for non response and reQTLs across all conditions. The analysis shows that reQTLs (purple) have a significantly lower replication rate based on π1 compared to non response eQTLs (green), with a Wilcoxon P-value = 4.9 x 10^-47^.

**Supplementary figure 7:**
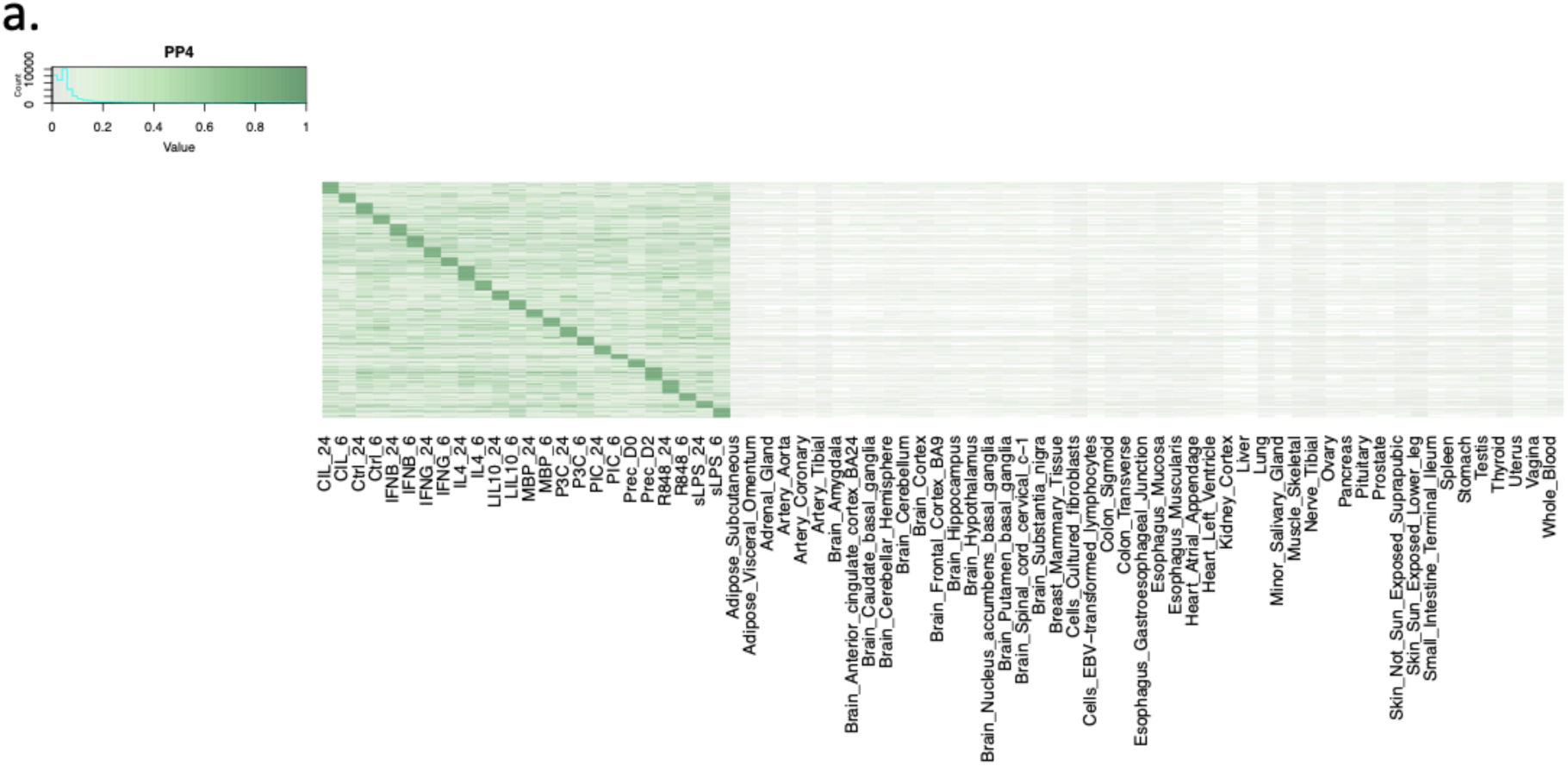
Colocalization evidence of eGenes with GWAS traits across GTEx tissues and MacroMap conditions. **a.** Heatmap showing the posterior probability of colocalization (PP4) for the 988 eGenes with higher colocalization evidence with one or more GWAS traits in MacroMap (PP4 >0.75) compared to GTEx (PP4 <0.5) across all GTEx tissues and conditions.

**Supplementary figure 8:**
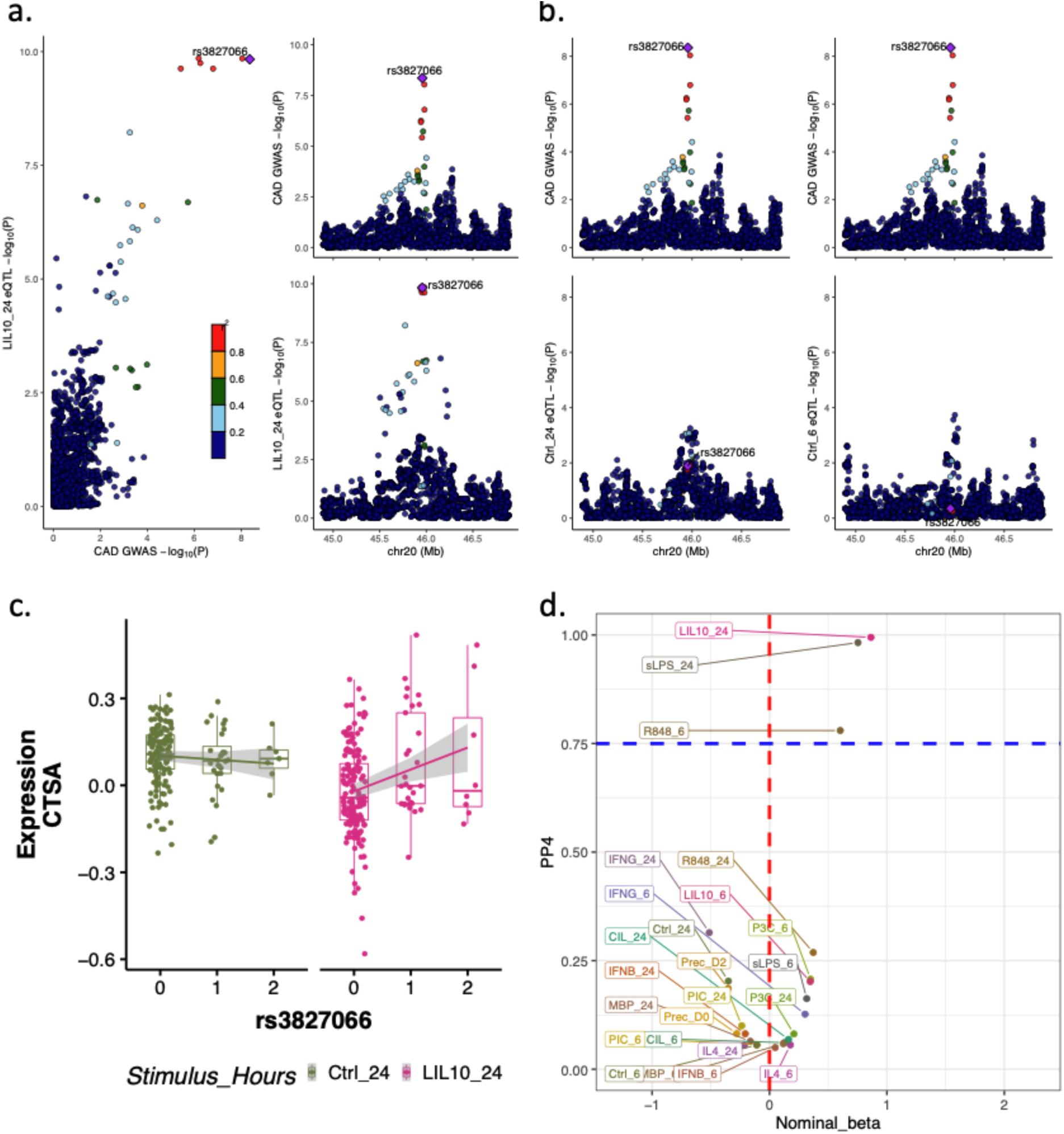
Colocalization between CAD GWAS hit and CTSA reQTL following LIL10 stimulation at 24h time point. **a.** Scatterplot and Manhattan plots of an apparent colocalization between CAD GWAS hit rs3827066 and CTSA reQTL after stimulation with LIL10 at the 24h time point, with the purple diamond representing the lead GWAS variant. **b.** Manhattan plots of the GWAS hit rs3827066 and both naive conditions (Ctrl at 24h/6h time point) where there is no detectable eQTL effect. **c.** rs3827066 is a reQTL following stimulation with LIL10_24 with higher expression for individuals carrying the alternative genotype (0=CC,1=CT,2=TT). **d.** Scatterplot which depicts the nominal betas (x-axis) (CTSA eQTL analysis in all conditions) and posterior probability of colocalization (PP4,y-axis) for the GWAS CAD variant (rs3827066) and CTSA eQTL summary statistics. Conditions with higher nominal betas show higher colocalization evidence (PP4).

**Supplementary figure 9:**
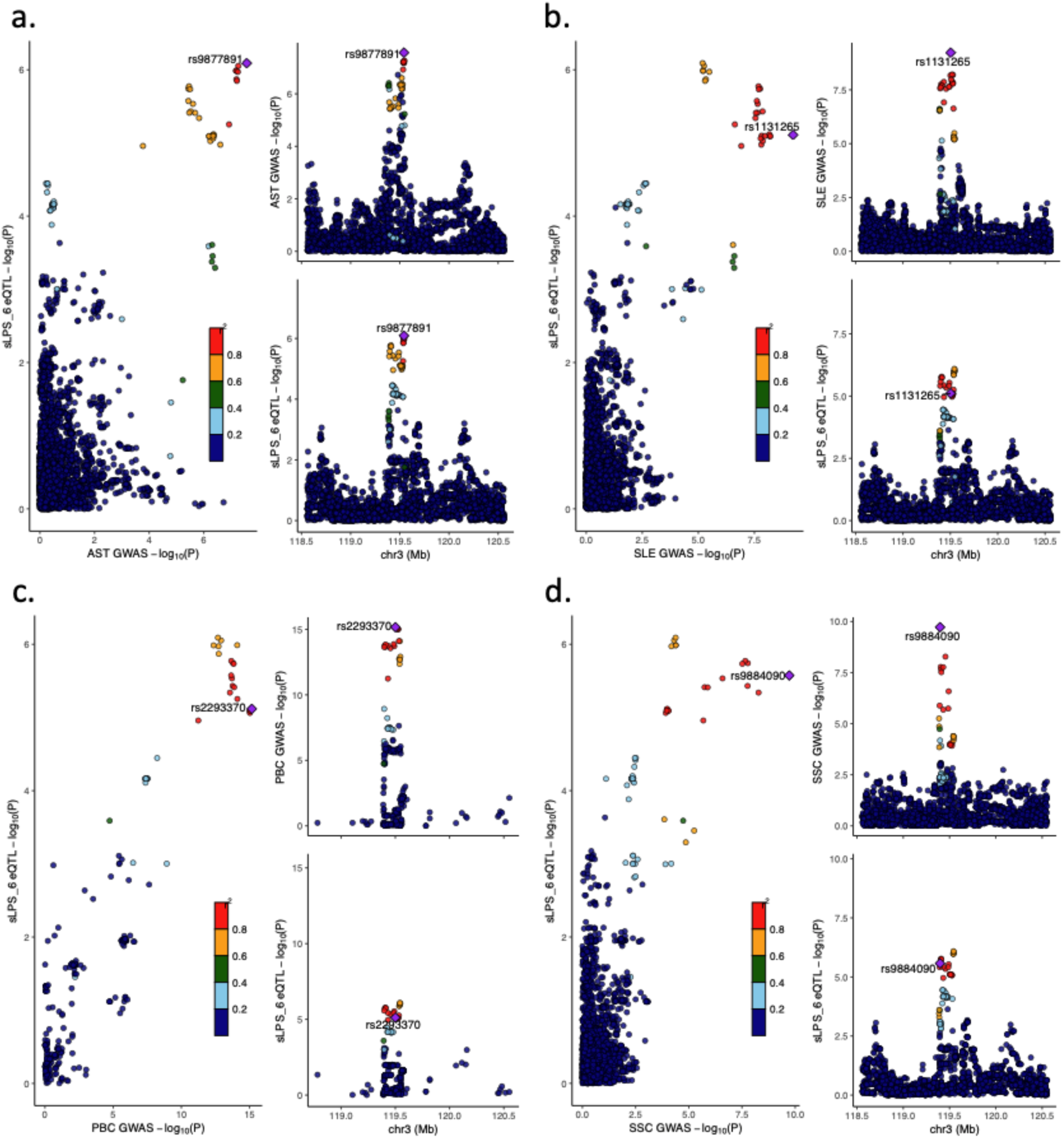
Colocalization of CD80 eQTL with GWAS hits in autoimmune diseases. **a-d.** Scatterplot and Manhattan plots demonstrate the colocalization between CD80 eQTL and four different GWAS studies in autoimmune diseases, namely **(a)** AST/Asthma (rs9877891,PP4=0.93), **(b)** SLE/Systemic Lupus Erythematosus (rs1131265,PP4=0.88) **(c)** PBC/Primary biliary_cirrhosis (rs2293370,PP4=0.91) and **(d)** SSC/Systemic Scleroderma (rs9884090,PP4=0.93) all following stimulation with sLPS at the 6h time point.

**Supplementary figure 10:**
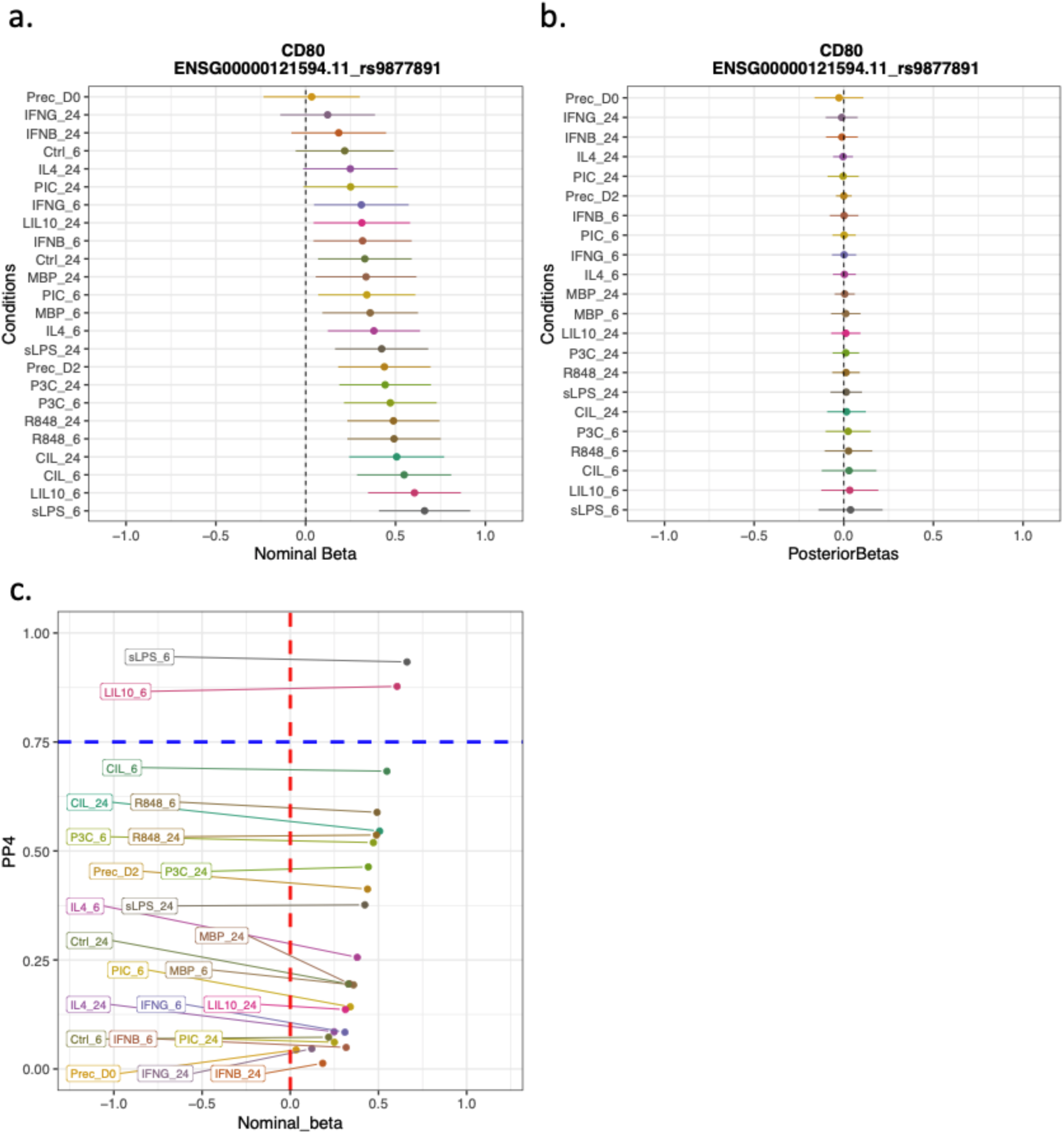
Exploring the potential of rs9877891 as a reQTL using condition-by-condition eQTL analysis and colocalization evidence. **a.** Metaplot of nominal betas and variances for all conditions in the condition-by-condition eQTL analysis. Specifically, it displays the results for the lead eQTL variant (rs9877891) in the sLPS_6 condition for CD80, which is also the lead GWAS variant for AST/Asthma (as shown in Supplementary Figure 9a). **b**.Metaplot which shows that mash analysis did not detect rs9877891 as a reQTL. Mash estimates the extent to which the eQTL effect size in each stimulated condition deviates from that of the baseline condition (Ctrl_24) and shrinks aggressively all effects towards zero. As a result, the effect size estimates for rs9877891 are very small. **c.** Scatterplot of the nominal betas (x-axis) (condition-by-condition eQTL analysis, all conditions) for the lead eQTL variant (rs9877891, in the sLPS_6 condition) and posterior probability of colocalization (PP4,y-axis) with AST GWAS hit. Conditions with slightly higher betas (sLPS_6, LIL10_6) compared to other conditions show strong colocalization evidence suggesting that the variant might be a true reQTL which remains undetected based on mash analysis.

